# Mechanism of cell polarisation and first lineage segregation in the human embryo

**DOI:** 10.1101/2020.09.23.310680

**Authors:** Meng Zhu, Marta N. Shahbazi, Angel Martin, Chuanxin Zhang, Berna Sozen, Mate Borsos, Rachel S. Mandelbaum, Richard J. Paulson, Matteo A. Mole, Marga Esbert, Richard T. Scott, Alison Campbell, Simon Fishel, Viviana Gradinaru, Han Zhao, Keliang Wu, Zijiang Chen, Emre Seli, Maria J. de los Santos, Magdalena Zernicka-Goetz

## Abstract

The formation of differential cell lineages in the mammalian blastocyst from the totipotent zygote is crucial for implantation and the success of the whole pregnancy. The first lineage segregation generates the polarised trophectoderm (TE) tissue, which forms the placenta, and the apolar inner cell mass (ICM), which mainly gives rise to all foetal tissues and also the yolk sac^1–3^. The mechanism underlying this cell fate segregation has been extensively studied in the mouse embryo^4,5^. However, when and how it takes place in the human embryo remains unclear. Here, using time-lapse imaging and 325 surplus human embryos, we provide a detailed characterisation of morphological events and transcription factor expression and localisation to understand how they lead to the first lineage segregation in human embryogenesis. We show that the first lineage segregation of the human embryo is triggered by cell polarisation that occurs at the 8-cell stage in two sequential steps. In the first step, F-actin becomes apically polarised concomitantly with embryo compaction. In the second step, the Par complex becomes polarised to form the apical cellular domain. Mechanistically, we show that activation of Phospholipase C (PLC) triggers actin polarisation and is therefore essential for apical domain formation, as is the case in mouse embryos^6^. Finally, we show that, in contrast to the mouse embryo, the key extra-embryonic determinant GATA3^7,8^ is expressed not only in extra-embryonic lineage precursors upon blastocyst formation. However, the cell polarity machinery enhances the expression and nuclear accumulation of GATA3. In summary, our results demonstrate for the first time that cell polarisation reinforces the first lineage segregation in the human embryo.

---

Studies in the mouse embryo demonstrate that the first lineage segregation begins with embryo compaction and polarisation on the third day of development (day 2.5) at the late 8-cell stage. The processes of compaction and polarisation lead to the closer apposition of neighbouring cells and the formation of an apical domain that orients towards the outside of the embryo^9–11^. The differential inheritance of the apical domain during the successive cell divisions generates polarised and non-polarised cells that sort to the outside and inside of the embryo, respectively^12–14^. Subsequently, cell polarity coordinates with cell position to control the expression of lineage-specific transcription factors, allowing the outside polarised cells to adopt TE identity and inner non-polarised cells to specify ICM fate^5,15^. Neither the timing of embryo polarisation, the sequence of its events, its mechanism, nor how it affects subsequent cell fate decisions in human embryos are known.

To address these fundamental questions, we first wished to determine the exact timing of cell polarisation in relation to compaction in the human embryo. Previous studies of human embryo polarisation were limited to the description of gross morphological features and the characterization of microscopic cellular structures by bright field imaging. These studies suggested that the cell-contact free membranes of human blastomeres become re-arranged between embryonic days 3 and 4, concurrent with the commencement of embryo compaction^16–18^. To address whether these morphological rearrangements relate to cell polarisation, we used surplus, *in vitro*-fertilised, cleavage-stage human embryos cryopreserved at embryonic day 3 and cultured them until embryonic day 4 for analysis by time-lapse imaging and immunofluorescence (Fig. 1a). We first carried out morphokinetic analyses on a total of 260 vitrified day 3 human embryos that at the start of the experiment had a mean number of 8 cells - 7.97±1.22 total cells and 7.67 ± 1.59 intact (non-degenerated) cells that increased to approximately 12 cells −12.23 ± 3.07 total cells and 12.14 ± 3.05 intact cells-after 24 hours in culture (Extended Data Fig. 1a-d). Interestingly, we found that the cell cycle time decreased from the 4/8-cell stage transition (22.15 hours ± 0.9192) to the 8/16-cell stage transition (14.49 hours ± 4.189) (Extended Data Fig. 1e). The rate of cell degeneration was very low, with 89.4% of the embryos showing no signs of degenerated cells after warming and 95.2% of the embryos showing no degenerated cells after 24 hours of culture (N=85, Extended Data Fig. 1f, g), indicating that the high quality of the embryos was not compromised after embryo warming. To provide an objective assessment of the dynamics of compaction, we measured the inter-blastomere angle between adjacent blastomeres in a representative middle plane through the embryo. When this angle increased to greater than 120°, we concluded that cells had compacted, in agreement with previous reports^6^. We found that the average cell number at the start of compaction was 10.14 ± 2.5 (N=83), increasing to 11.57 ± 3.1 (N=54) by completion of compaction (Extended Data Fig. 1h). In addition, we analysed compacting embryos in which the exact times of fertilisation and cryopreservation were known. This revealed that the completion of compaction was accomplished at 80.2 ± 8.5 hours post-fertilisation (h.p.f.) (Extended Data Fig. 1i-k). These observations indicate that the onset of compaction in human embryos is heterogeneous, and while it starts at the 8-cell stage, it extends to the 8/16-cell stage. This is consistent with previous observations^16,17,19^ and contrasts with the timing of compaction in the mouse embryo where this process is completed by the end of the 8-cell stage^6,20^. Hence, the process of compaction in the human embryo is prolonged in comparison to the mouse embryo.

**Figure 1.**
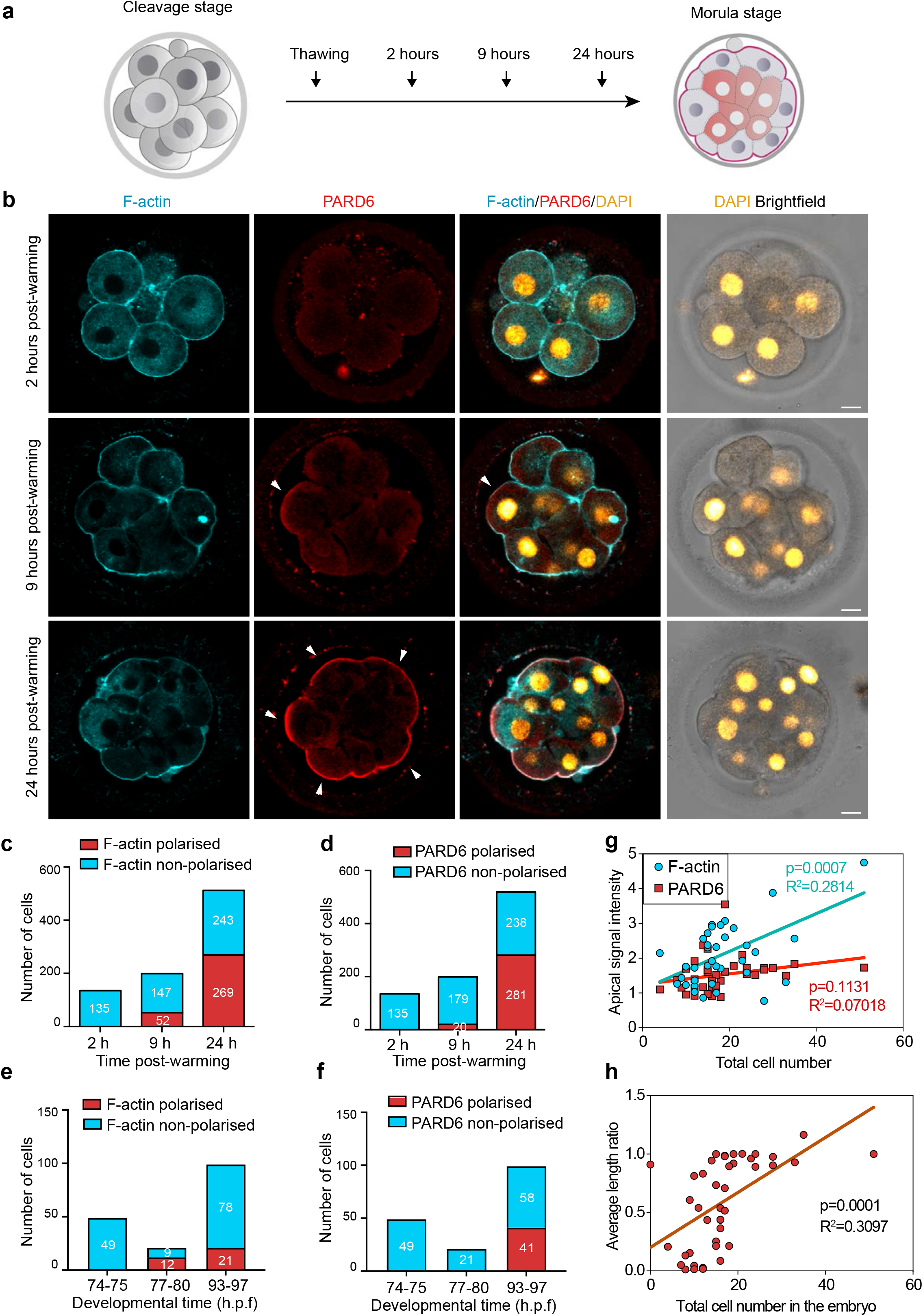
Timing of compaction and polarisation in human embryos. **a,** Scheme for human embryo culture. Surplus *in vitro* fertilised human embryos were warmed at day 3, and cultured for 2, 9 or 24 hours(h) to examine the localisation of polarisation markers. **b,** Representative images of human embryos fixed at different developmental time-points (as shown in a) and immunostained for F-actin and PARD6b. Arrowheads indicate the apical domain. **c,** Quantification of number of cells showing polarised or non-polarised F-actin in different developmental time-points. **d,** Quantification of number of cells showing polarised or non-polarised PARD6. For both c and d, the number in each bar represents the number of cells analysed. N=28 embryos for 2 h post-warming; N=21 embryos for 9 h post-warming and N=65 embryos for 24 h post-warming. N=9 independent experiments. **e,** Quantification of the number of cells showing polarised or non-polarised F-actin in different post-fertilisation time-points. **f,** Quantification of the number of cells showing polarised or non-polarised PARD6 in different post-fertilisation time-points. For both e and f, the number in each bar represents the number of cells analysed. N=6 74-75 h.p.f., N=2 77-78 h.p.f. and N=9 h.p.f. embryos. **g,** Correlation between F-actin/PARD6 apical enrichment and cell numbers. Apical enrichment of F-actin and PARD6 is measured as the ratio of signal intensity on the apical surface to the cell-cell contacts. Apical enrichment of F-actin and PARD6 increases as embryo cell numbers increase. **h,** Correlation between the length of the apical domain (based on the PARD6 immunostaining) and embryo cell numbers. The length of the apical domain increases as the embryo cell number increase. Each dot represents one analysed cell. h.p.f.: hours post-fertilisation. 7 independent experiments. Scale bars, 15 μm.

We next wished to determine the sequence of developmental events ensuring human embryo polarisation in relation to compaction. In the mouse embryo, the polarisation of the Par complex to the apical domain serves as the crucial developmental transition that control the expression of TE lineage specific transcription factors^4^. Transcriptomic analyses indicate that Par complex components, such as PARD6, are also expressed in the human embryo at the 8-cell stage^7,21^. We therefore set out to determine the localisation of PARD6 during embryo compaction. Our recent studies in the mouse embryo revealed that the Par complex acquires apical localisation following two distinct steps. An initial apical enrichment of F-actin is concomitant with embryo compaction followed by the polarisation of the Par complex to the apical surface^6^. To determine whether this localisation pattern also occurs in the human embryo, we fixed embryos at three different time points during *in vitro* culture (2, 9 or 24 hours) and examined the localisation of PARD6 and F-actin (Fig. 1a). We found that F-actin localised circumferentially around the cell cortex and became apically enriched upon compaction. None of the embryos were compacted after 2 hours of culture in agreement with our time-lapse studies (N=18 embryos). At this stage, both cortical Factin and PARD6 were uniformly distributed within the embryos, indicating that they had not yet initiated polarisation (N=135 cells; 18 embryos) (Fig. 1b-d). However, 7 hours later 26% of embryos had become fully compacted (N=199 cells; 19 embryos). All compacted cells displayed an apical enrichment of cortical F-actin at the cell-contact free surface (Fig. 1b-d). In contrast, only half of these compacted cells showed PARD6 polarisation (26.3%, N=199 cells) (Fig. 1b-d). The proportion of compacted embryos in which both F-actin and PARD6 were polarised increased significantly during the next 15 hours of culture (91.0% for compaction, N=11 embryos; 71.5% for PARD6 polarisation, N=628 cells) (Fig. 1b-d). Quantification of cell polarisation in embryos in which the exact times of fertilisation and cryopreservation were known revealed a similar pattern; none of the embryos showed F-actin nor PARD6 polarisation (N=49 cells; 6 embryos) at 74-75 h.p.f; F-actin polarisation was observed at 77-80 h.p.f. (57.1%; N=21 cells; 2 embryos), while PARD6 polarisation was detected 93-97 h.p.f. (41.4%; N=99 cells; 9 embryos) (Fig. 1e-f). The level of PARD6 apical enrichment, as well as the size of the PARD6 domain, showed a positive correlation with embryo cell numbers, indicating that the apical localisation of PARD6 increases as embryos develop (Fig. 1g-h). We also confirmed that other components of the apical Par polarity complex such as atypical protein Kinase C (aPKC) localises to the cell-contact free surface at the same developmental time frame (N=5, Extended data Fig. 2a). Globally, these results indicate that polarisation of the human embryo follows two steps: in the first, F-actin becomes polarised concomitantly with embryo compaction, and in the second, the Par complex becomes polarised.

We next wished to determine the mechanism of human embryo polarisation. Our recent studies in the mouse embryo showed that polarisation is regulated by the PLC-Protein Kinase C (PKC) pathway, which enables the recruitment of the actin-myosin complex to the cell membrane^6^. We therefore analysed whether the PLC-PKC pathway is also responsible for polarisation of the human embryo. To address this question, we first cultured human embryos from day 3 in the presence of the PLC inhibitor U73122^22^ (Fig. 2a) using three different concentrations (5 μM, 7.5 μM and 10 μM), and analysed the localisation of F-actin and PARD6. At the lowest concentrations (5 μM), U73122 failed to have an impact on cell polarisation (N=42 control embryos; N=19 DMSO-treated embryos; N=25 U73122-treated embryos), whereas at a higher concentration (7.5 μM) U73122 significantly reduced the proportion of polarised cells and the level of apical enrichment for PARD6 (N=42 control embryos; N=24 DMSO-treated embryos; N=24 U73122-treated embryos) (Fig. 2b-f and Extended Data Fig. 2b and 3a, b), but without an obvious effect on embryo cell number (Fig. 2e), suggesting a specific effect of U73122 on cell polarisation. By contrast, high levels of PLC inhibitor (10 μM) affected embryo development and led to apoptosis (N=66 control embryos; N=23 DMSO 5 μM-treated embryos; N=31 U73122 5 μM-treated embryos; N=20 DMSO 7.5 μM-treated embryos; N=20 U73122 7.5 μM-treated embryos; N=9 DMSO 10 μM-treated embryos; N=10 U73122 10 μM-treated embryos, Extended Data Fig. 3c). Based on these results, we carried out subsequent experiments using low concentrations of U73122 (5 μM and 7.5 μM). Next, we analysed whether the PLC pathway controls compaction. We observed a mild and non-statistically significant effect of PLC inhibition on the percentage of embryos that compacted during the first 24 hours of culture (N=66 control embryos; N=23 DMSO 5 μM-treated embryos; N=31 U73122 5 μM-treated embryos; N=27 DMSO 7.5 μM-treated embryos; N=27 U73122 7.5 μM-treated embryos Extended Data Fig. 3d). These results indicate that PLC is a major regulator of cell polarisation, but not of compaction.

**Figure 2.**
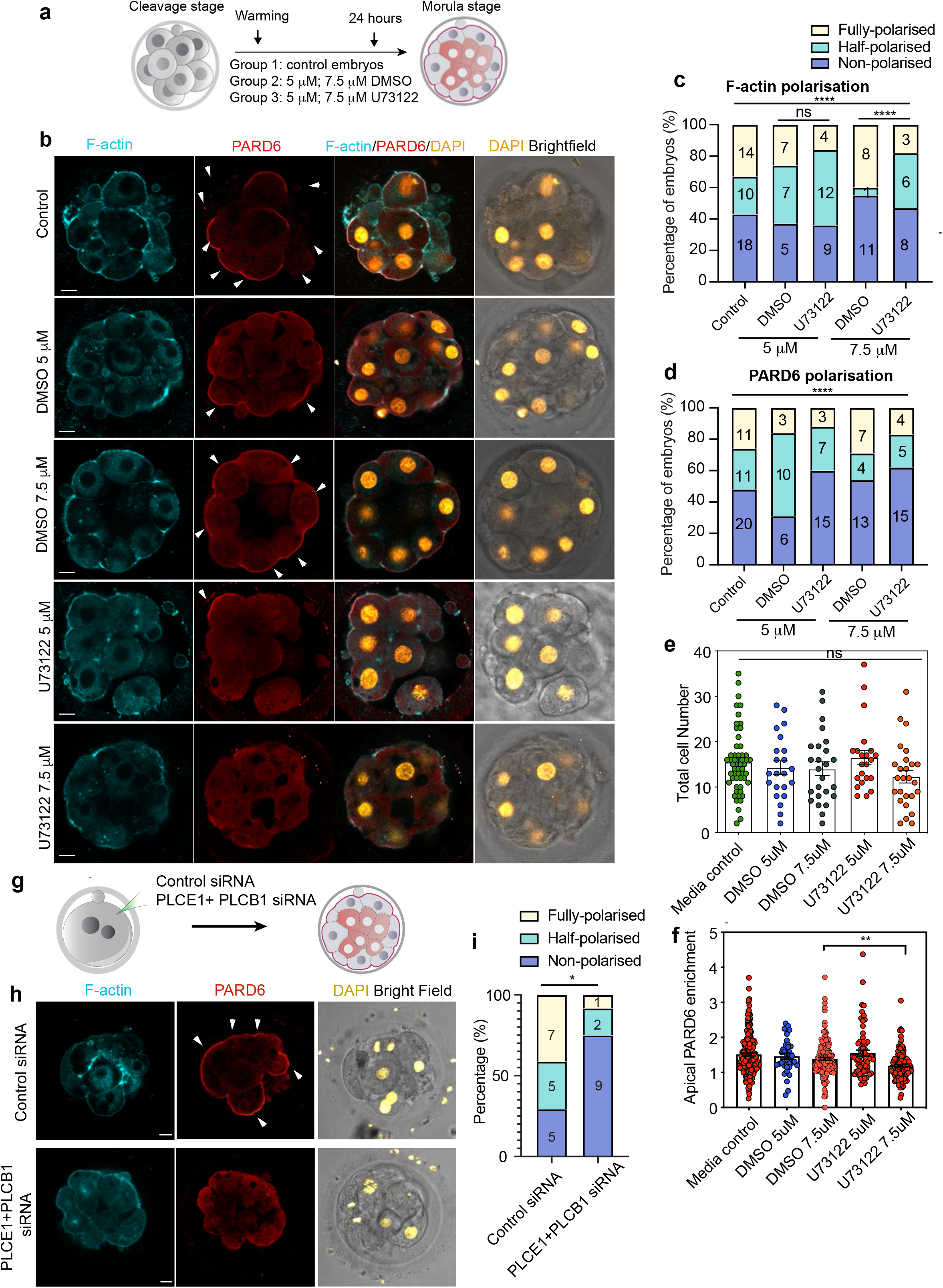
PLC activity regulates cell polarisation in the human embryo. **a,** Scheme for PLC inhibitor treatment. **b,** Representative images of *in vitro* fertilised human embryos warmed at day 3, cultured for 24 h (see scheme in Fig. 1a) and treated with or without DMSO or U73122 at different concentrations to reveal the localisation of F-actin and PARD6. Arrowheads indicate the apical domain. **c-d**, Quantification of the percentage of embryos showing F-actin (c) and PARD6 (d) polarised cells in embryos from panel b. Embryos were binned into three different categories based on the ratio of polarised cells to total cell numbers as shown in Extended Data Fig. 3a. ns:non-significant, ****p<0.0001. Chi-Square test. N=42 control embryos, N=19 DMSO 5μM embryos, N=25 U73122 5 μM embryos, N=20 DMSO 7.5μM embryos and N=17 U73122 7.5 μM embryos. **e,** Quantification of the total number of cells in embryos of panel b. Each dot represents one embryo. N= 55 embryos for media control group, N=22 embryos for DMSO 5μM, N=25 embryos for DMSO 7.5μM, N=22 embryos for U73122 5μM and N=26 embryos for U73122 7.5μM. Data is shown as mean ± S.E.M. ns: non-significant, Kruskal-Wallis test. **f,** Quantification of apical PARD6 fluorescence intensity in embryos from panel b. N=204 cells from 38 embryos for Media control; N=41 cells from 10 embryos for DMSO 5μM; N=158 cells from 17 embryos for DMSO 7.5μM, N=72 cells from 14 embryos U73122 5μM and N=126 cells from 24 embryos U73122 7.5μM. Each dot represents one analysed cell. ***p<0.001, Kruskal-Wallis test with a multiple comparisons test. Data is shown as mean ±S.E.M. **g**, Scheme for human zygote injection of PLCE1/PLCB1 siRNA. **h**, Representative images for embryos injected with control siRNA or PLCE1/PLCB1 siRNA and cultured until embryonic day 4 to reveal localisation of F-actin, PARD6, and DAPI. **i,** Quantification of the percentage of embryos showing PARD6 polarised cells in embryos from panel h. Number in each bar indicates the number of embryos analysed. N=18 embryos for control siRNA injected group; N = 12 for PLCB1 + PLCE1 siRNA injected group. * p<0.05. Fisher’s exact test. 5 independent experiments for PLC inhibitory treatments and 3 independent experiments for PLCE1/PLCB1 siRNA injections. Scale bars, 15μm.

To substantiate the role of PLC in cell polarity establishment of the human embryo, we next used RNA interference (RNAi)^23^ to down-regulate PLC expression at the cleavage stage and determine its effects on embryo polarisation. Specifically, we chose to deplete PLCE1 and PLCB1, as the level of their mRNA expression rank the highest among all functional PLC isoforms in human embryos between the 2-to the 8-cell stage (Extended Data Fig. 3e)^24^. To validate the siRNAs, we either injected the siRNA into human embryos or transfected them into HeLa and 293T cells. These experiments revealed a 70% downregulation of PLCE1 and PLCB1 (Extended Data Fig. 3f). Next, we injected control and PLCE1/PLCB1 siRNAs into human embryos at the zygote stage and allowed them to develop until day 4 to analyse the localisation of PARD6. The analysis of PARD6 localisation by immunofluorescence revealed a significantly reduced number of polarised cells in PLCE1/PLCB1 siRNA embryos (Fig. 2g-i; N=18 embryos for control siRNA embryos; N=12 embryos for PLCB1+PLCE1 siRNA embryos), in agreement with the result of PLC inhibitor treatment. Thus, both the pharmacological and RNAi approach indicate that PLC activity regulates cell polarisation during early human embryo development.

Finally, we wished to determine whether embryo polarisation is critical for initiation of cell fate specification. To address this question, we analysed the spatiotemporal expression of the transcription factor GATA3, which is a key marker of TE specification and based on single-cell RNA sequencing is already induced at the compacting morula stage^7^. We found that embryos lacked GATA3 expression after either 2 or 9 hours in culture, regardless of whether individual cells within the embryo were polarised or not (Fig. 3a-b, N=135 cells; 18 embryos for 2 hours and N=199 cells; 19 embryos for 9 hours). In contrast, we detected GATA3 nuclear localisation in 53.5% of the cells after 24 hours of culture, and therefore on day 4 (Fig. 3a-b, N=519 cells; 75 embryos). Strikingly, GATA3 was localised to the nucleus in both polarised and non-polarised cells, although the nuclear signal intensity of GATA3 was significantly higher in polarised cells. These results suggest that in human embryos GATA3 expression is initiated independently of embryo polarisation, but that its expression and nuclear localisation is reinforced by the acquisition of apicobasal polarity (Fig. 3a, c). To understand whether cell polarisation also promotes nuclear localisation of other TE transcription factors, we also examined the localisation of YAP1 in inner and outer polarised cells. We found that YAP1 was also significantly enriched in outer polarised cells in comparison to inner apolar cells (N=4 embryos; Extended Fig. 4a-b). These results suggest that cell polarisation promotes the accumulation of TE-associated transcription factors in the nucleus as early as the morula stage, similar to what happens in the mouse embryo.

**Figure 3.**
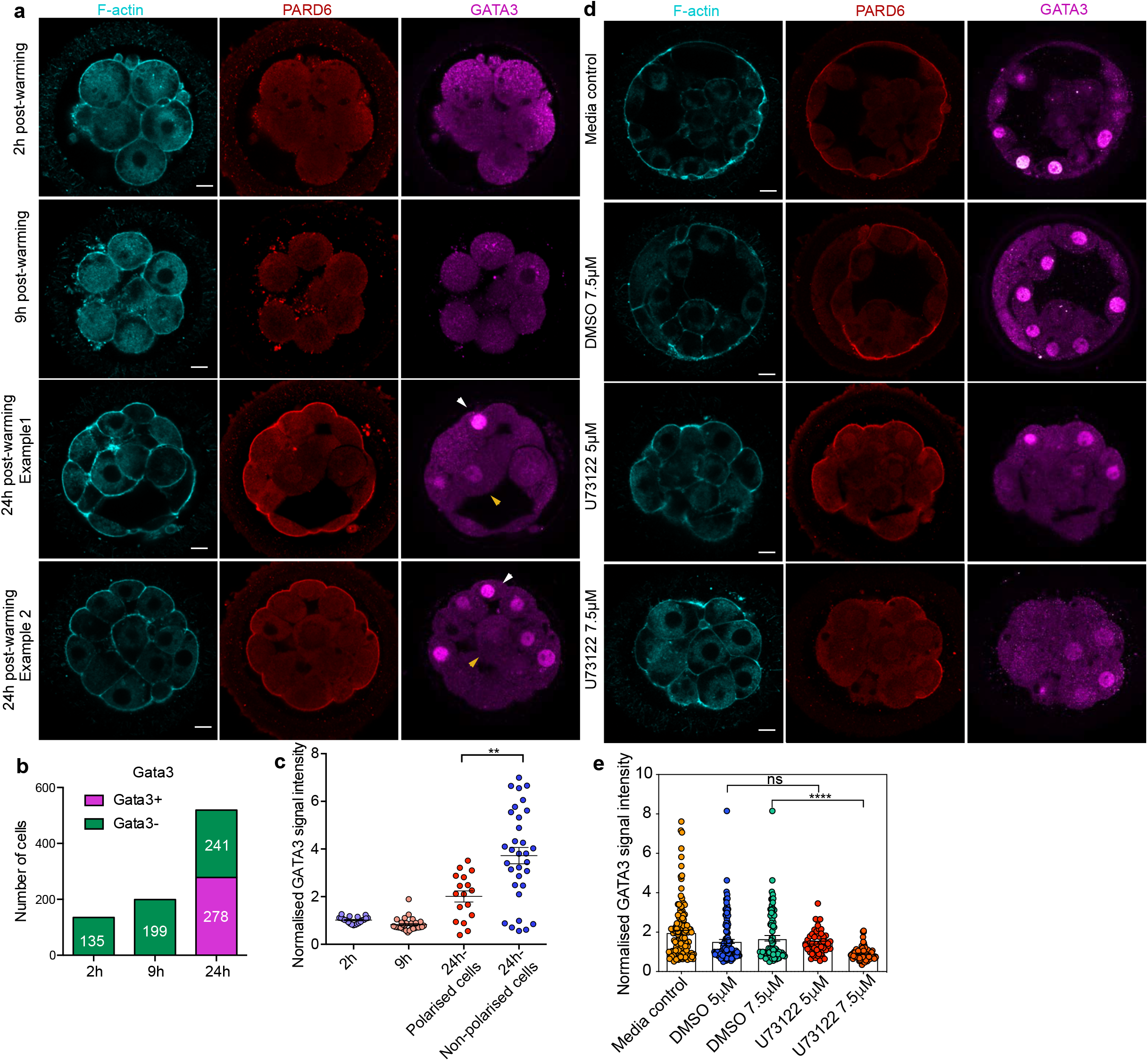
GATA3 expression is initiated independently of cell polarisation. **a,** Representative images of *in vitro* fertilised human embryos warmed at day 3 and cultured for 2, 9 or 24 h (see scheme in Fig. 1a) to reveal the localisation of F-actin, PARD6 and GATA3. GATA3 expression was undetectable until 24 h post-warming. White arrowheads indicate outer cells; yellow arrowheads indicate inner cells. Inner cells also express GATA3 albeit at lower levels. **b,** Quantification of the number of GATA3 positive cells in embryos from panel a. Cells that have higher nuclear GATA3 signal comparing to cytoplasm are categorised as GATA3+ cells. Numbers in each bar indicates the number of cells analysed. N=17 embryos for 2 h N=20 embryos for 9 h and N=38 embryos for 24 h. **c,** Quantification of GATA3 nuclear signal in embryos from panel a. Each dot represents one cell. Data shown as mean ±S.E.M. The GATA3 nuclear signal has been calculated as the nucleus to cytoplasmic signal ratio in each cell. N=5 embryos for 2 h, N=5 embryos for 9 h and N=5 embryos for 24 h. **p<0.01; student’s t test. **d,** Representative images of embryos treated with different concentrations of U73122, cultured from day 3 to day 4 and stained for F-actin, PARD6 and GATA3. **e,** Quantification of GATA3 nuclear signal intensity in embryos from panel d. N=142 cells from 18 embryos for Media control; N=37 cells from 6 embryos for DMSO 5 μM; N=84 cells from 12 embryos for DMSO 7.5 μM, N=56 cells from N=9 embryos U73122 5 μM and N=90 cells from N=13 embryos U73122 7.5 μM. Each dot represents one cell analysed. ns: non-significant. **** p < 0.0001, Kruskal-Wallis test. 2 independent experiments (a-c) and 5 independent experiments (d-e). Scale bars, 15 μm.

To further confirm the role of cell polarisation in the expression of TE transcription factors, we wished to analyse GATA3 expression after U73122 treatment, as this would inhibit PLC and therefore embryo polarisation. To this end, we applied U73122 at 5 μM and 7.5 μM, cultured the embryos from day 3 to 4 and analysed GATA3 expression. We found that U73122 treatment led to a significant reduction of the nuclear GATA3 signal intensity (Fig. 3d-e, N=142 cells for control embryo; N=37 cells for DMSO 5 μM treated group; N = 84 cells for DMSO 7.5 μM group; N=56 cells for U73122 5 μM treated group; N = 90 cells for U73122 7.5 μM treated group), but it did not affect the number of GATA3 positive cells per embryo (Extended Data Fig. 4c, N=46 control embryos; N=19 DMSO 5 μM-treated embryos; N=27 U73122 5 μM-treated embryos; N=24 DMSO 7.5 μM-treated embryos; and N=23 U73122 7.5 μM-treated embryos). Together, these results demonstrate that the onset of expression of the critical TE determinant, GATA3, is controlled by a polarity-independent pathway, but the levels of nuclear GATA3 are reinforced by a polarity-dependent pathway^25^.

The fact that expression of the TE lineage determinant, GATA3, is initiated independently of embryo polarisation led us to examine when inside cells, precursors of the ICM, are first generated. In the mouse embryo, inside cells are produced after all blastomeres become polarised at the 8-cell stage and by subsequent cell divisions from the 8-cell to 16-cell stage^4^. Consequently, inner cells can only be observed in embryos having more than eight polarised cells, but whether the same is true in the human embryo remains unknown. To address this question, we first analysed the number of inside cells and their correlation with cell polarity in relation to overall cell numbers in fixed human embryos. Surprisingly, we found that in 37% of embryos, inside cells were present with less than eight outer polarised cells (Fig. 4a-b, N=27 embryos). This suggests that the generation of inside cells in the human embryo might not require the preceding establishment of embryo polarisation. In addition, we found that the size of the inside cells was highly variable, ranging from 229.123 to 1896.4 μm^2^ (Fig. 4c-d). This partially overlapped with the size of 8-cell stage blastomeres, which ranges from 895.432 to 2627.03 μm^2^ (Fig. 4d). Specifically, we found that 34% (18 out of 53 embryos) of inside cells were bigger than the smallest 8-cell stage blastomere analysed. This suggests that cells can be internalised without undertaking asymmetric cell division at the 8-cell to 16-cell stage transition. To directly assess this possibility, we performed time-lapse imaging of live fluorescently labelled human embryos to record cell positions during the process of compaction. To be able to follow cells in live embryos, we labelled the cell membrane using a live-membrane dye (FM 4-64FX). From the 5 human embryos imaged, we found one embryo in which a cell became positioned to the inside compartment during the process of compaction (Extended Fig. 4d). Therefore, whereas polarisation and subsequent asymmetric cell divisions are critical to generate inside cells in mouse embryos, in human embryos inside cells can be generated independently of polarity and asymmetric cell divisions, revealing mechanistic differences of TE/ICM segregation between the two species.

**Figure 4.**
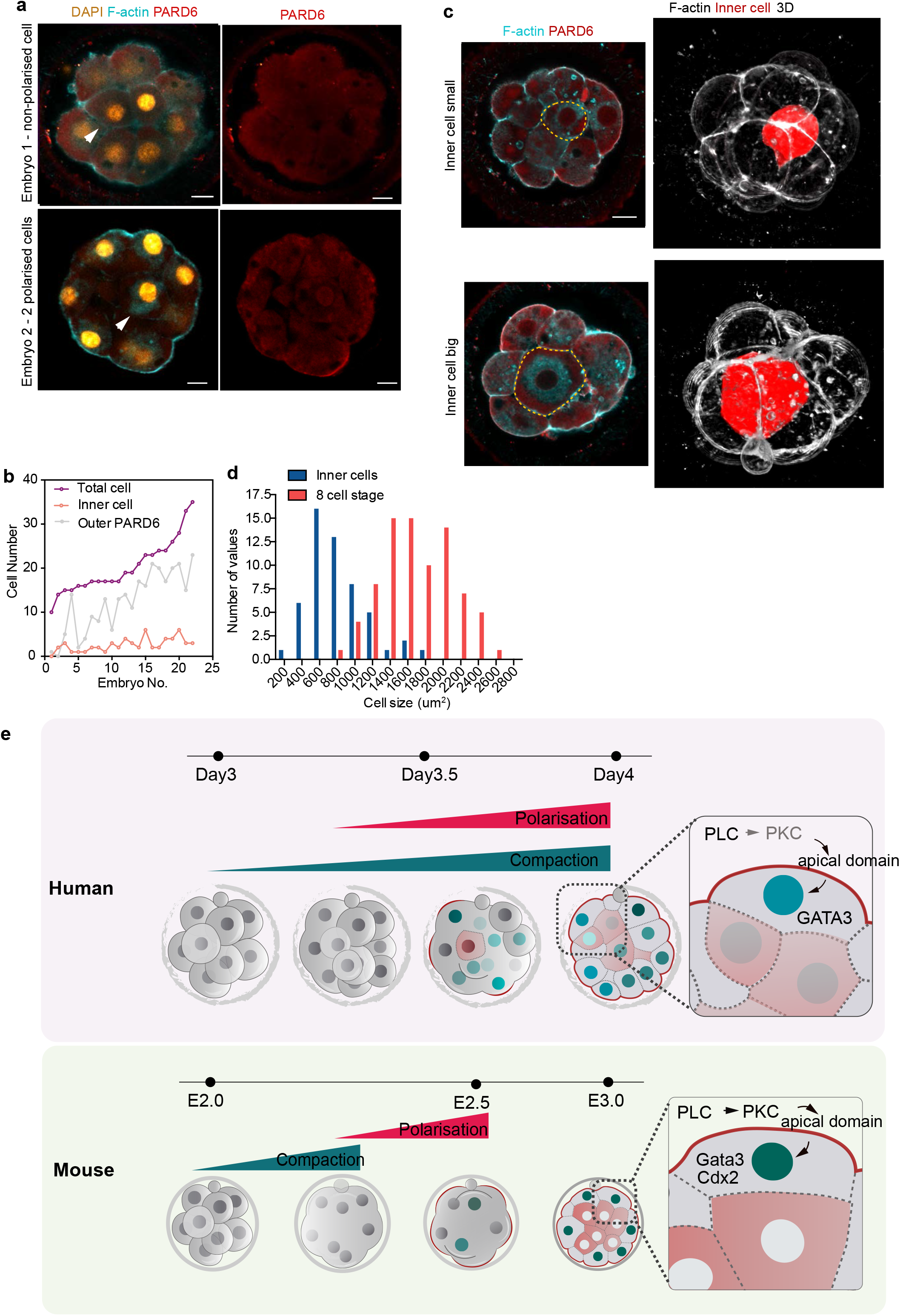
Inner cells in human embryo can be generated by a polarity independent mechanism. **a,** Representative images of embryos that have inner cells with low number of outer polarised cells. White arrowheads indicate the presence of inner cells. **b,** Line chart showing the relation between the number of inner cells, the number of PARD6 positive cells and total cell number in embryos that have inner cells. **c,** Representative images of embryos showing inner cells with different sizes. Dotted lines indicate the outline of the inner cells. **d,** Quantification of the distribution of the size of inner cells in comparison with the size of 8-cell stage blastomeres. N=53 inner cells from 21 embryos; and N=80 8-cell stage blastomeres from 10 embryos. **e,** Model summarising the main findings of this study. F-actin polarisation precedes Par complex polarisation and it is triggered by PLC activation. Blastomeres initiate expression of TE factors independently of the polarity machinery, but polarisation reinforces a TE fate. 5 independent experiments. Scale bars, 15 μm.

In summary, our results uncover the timing and the mechanism behind embryo polarisation and its function in the first lineage segregation in the human embryo (Fig. 4e). We show that the mechanism that triggers embryo polarisation is evolutionarily conserved between mouse and human. However, we also show that the onset of cell fate specification – cell allocation and expression of lineage determining genes – is independent of cell polarisation. Instead, cell polarisation is needed to reinforce a TE fate, which becomes established at the blastocyst stage^7^. These results provide important insight into human embryo development at a stage when a crucial morphogenetic transition occurs.

## Author contribution

M.Z., M.N.S., A.M. and M.J.S., designed and performed experiments and analysed the data. CX.Z. and M.B. performed microinjection experiments; M.A.M. performed an inhibitor treatment experiment. A.C., S.F., R.P. provided human embryos. E.S., R.T. S., M.N.S., M.J.S., R.S.M., Z-J.C. and H.Z. secured ethical approvals. B.S., M.E., K-L.W., helped with experiments. Z-J.C., M.J.S., R.T.S. provided financial support. V.G. provided funding and supervision to M.B. M.Z., M.N.S. and M.Z-G. wrote the manuscript. M.Z-G. conceived and supervised the project, helped with data analyses and interpretation, and provided funding.

## Acknowledgement

We are thankful to David Glover for comments on the manuscript. We thank Marta Perez Sanchez, Antonia Weberling, Bailey Weatherbee and Ali Ahmady for the help with human embryo culture. We are grateful to USC Fertility and HCLD for their support. M.Z. is funded by Leverhulme Trust. M.N.S. is funded by the European Molecular Biology Organization (EMBO) and the Medical Research Council (MRC, MC_UP_1201/24). Work in the laboratory of M.Z-G. on human embryos is funded by Wellcome Trust (207415/Z/17/Z), Open Philanthropy Grant, Curci and Weston Havens Foundations. Work in the laboratory of Zi-Jiang Chen is funded by The National Key Research and Development Program of China (2018YFC1004000) and Shandong Provincial Key Research and Development Program (2018YFJH0504).

## Extended Data Figure Legends

**Extended Data Figure 1:**
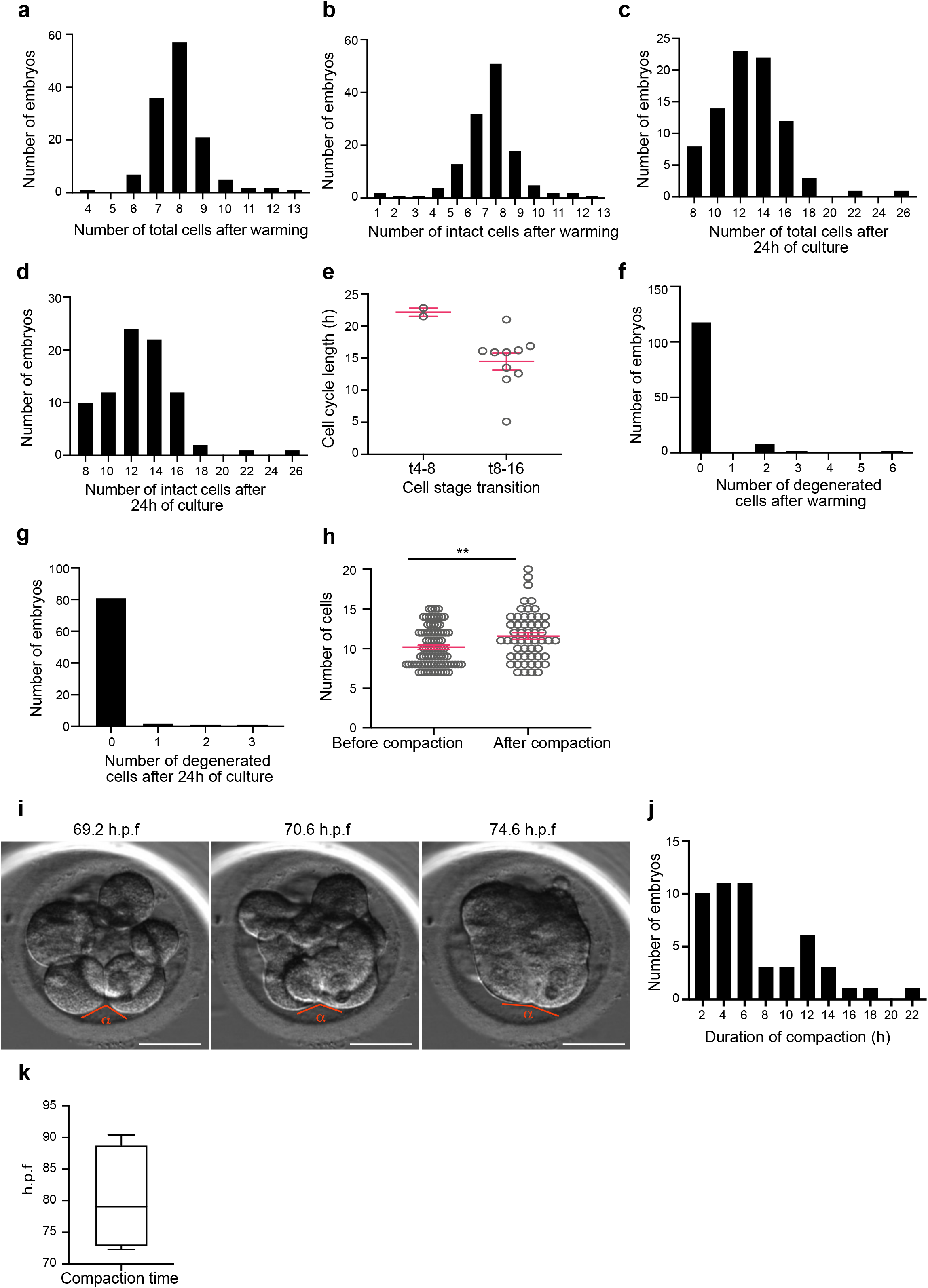
morpho-kinetic analysis of human embryos cultured *in vitro* from day 3 to day 4. **a-b,** Histograms showing the number of total (a) and intact (b) cells at the time of embryo warming, N=132 control embryos. **c-d,** Histograms showing the number of total (c) and intact (d) cells after 24 h of embryo culture, N=85 control embryos. **e,** Cell cycle length shown at the 4 to 8 (t4-8) and 8 to 16 (t8-16) cell stage transitions. Each dot represents an individual embryo. N=2 t4-8 and N=10 t8-16 control embryos. Data is shown as mean ± S.D. **f,** Histogram showing the number of degenerated cells at the time of embryo warming, N=132 control embryos. **g,** Histogram showing the number of degenerated cells during 24 h of embryo culture, N=85 control embryos. **h,** Number of cells before and after compaction. Each dot represents an individual embryo. Data is shown as mean ± S.E.M. N=83 (before) and N=54 (after) control embryos. **p=0.0083, Mann Whitney U test. **i,** Brightfield images of an embryo undergoing compaction. The time post-fertilisation is indicated in h as h.p.f. The angle measured to assess compaction is indicated. Scale bars, 10 μm. **j,** Histogram showing the time period between the initiation and the end of compaction. N=50 control embryos. **k,** Time at which compaction is completed in embryos with known fertilisation time. N=4 control embryos. Data is shown as a box and whiskers plot from the minimum to the maximum value. 2 independent experiments.

**Extended Data Figure 2:**
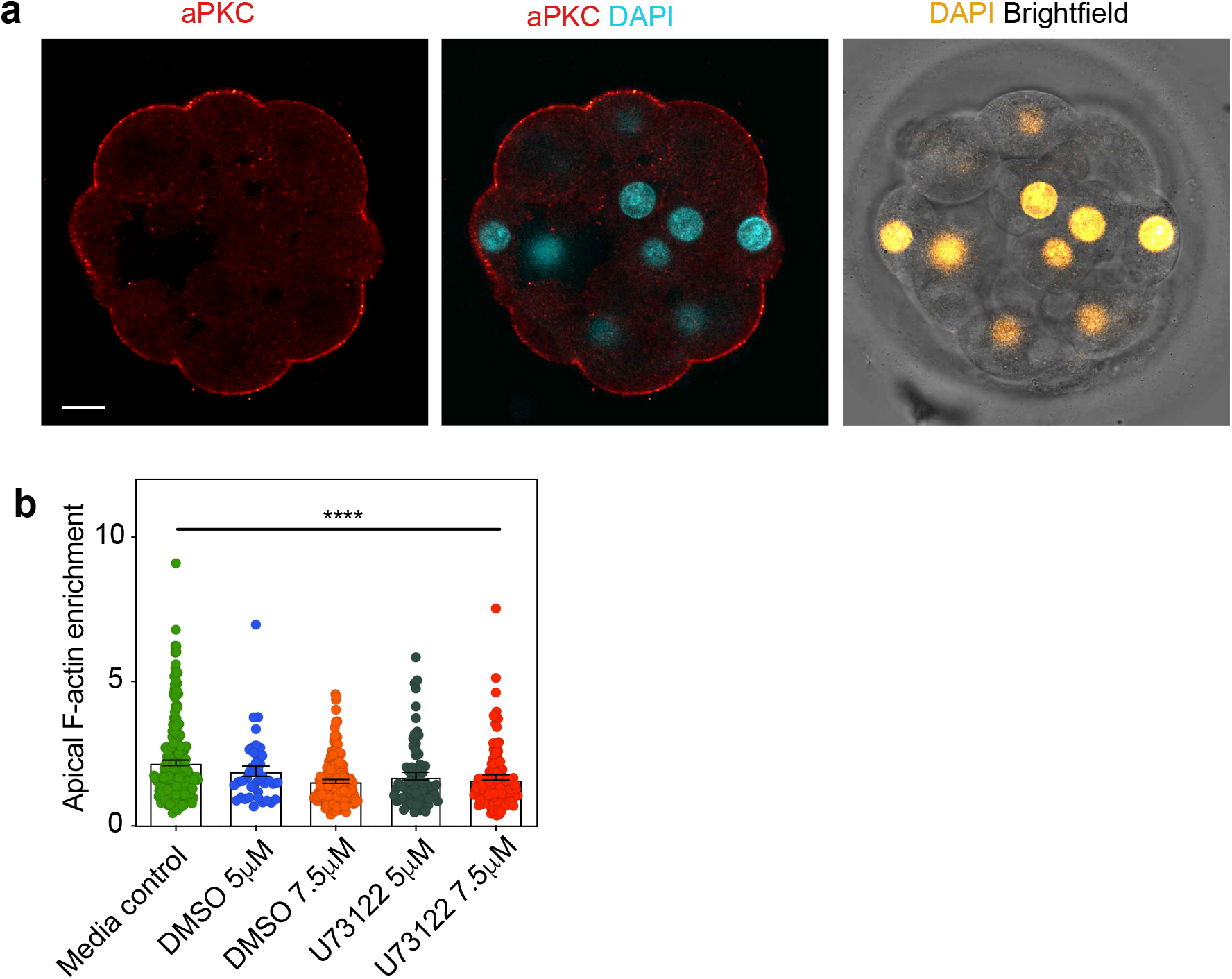
localisation of polarisation proteins at the morula stage in human embryo. **a,** Human embryos at embryonic day 4 were immunostained with aPKC, F-actin and DAPI. aPKC localises to cell-contact free surface in a similar fashion with PARD6.N=5 embryos. **b,** Quantification of apical F-actin fluorescence intensity in embryos from Fig. 2b. Each dot represents one analysed cell. N=204 cells from 38 embryos for Media control; N=41 cells from 10 embryos for DMSO 5μM; N=158 cells from 17 embryos for DMSO 7.5μM, N=72 cells from 14 embryos U73122 5μM and N=126 cells from 24 embryos U73122 7.5μM. ***p<0.001, Kruskal-Wallis test. Scale bars, 15 μm.

**Extended Data Figure 3:**
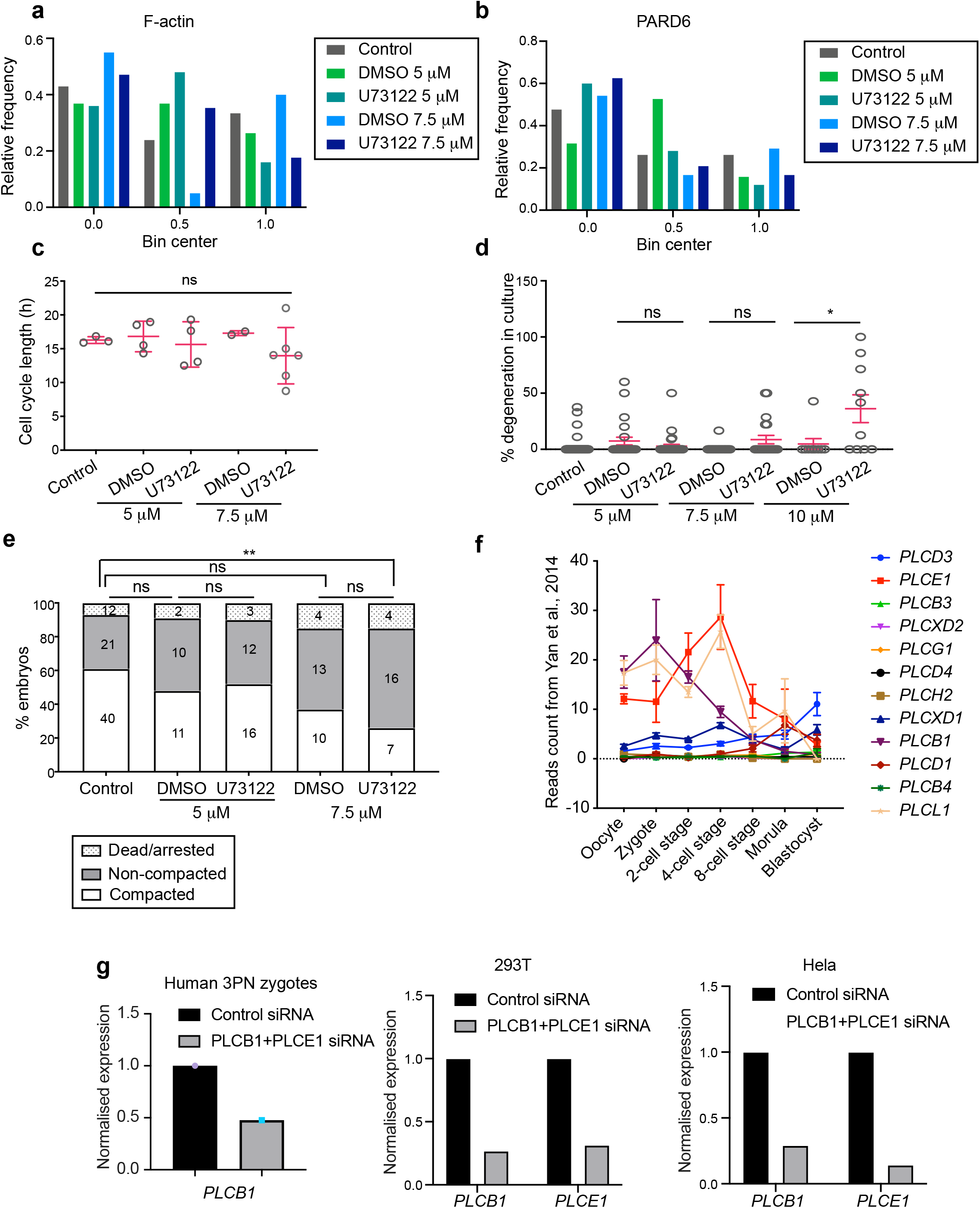
morpho-kinetic analysis of WT and PLC inhibitor treated embryos. **a-b,** Histograms showing the ratio of F-actin (a) and Par6 (b) polarised cells in embryos from Fig. 2b. Data was binned into three different groups. **c,** Percentage of degenerated cells in embryos from Fig. 2b. Each dot represents an individual embryo. Data is shown as mean ± S.D. N=66 embryos (control), N=23 embryos (DMSO 5 μM), N=31 embryos (U73122 5 μM), N=20 embryos (DMSO and U73122 7.5 μM), N=9 embryos (DMSO 10 μM) and N=10 embryos (U73122 10 μM). Kruskal-Wallis test with a multiple comparisons test. *p=0.0355, ns: non-significant. **d,** Compaction analysis in control and PLC inhibitor treated embryos. The specific number of embryos per category is indicated. Ns: non-significant, **p=0.0099. **e**, Expression profile for all PLC isoforms at different stages of human preimplantation development. Data retrieved from Yan et al., 2014. **f,** Expression level of PLCE1 and PLCB1 in human 3PN zygotes injected with control siRNA or PLCE1+ PLCB1 siRNA; or Hela and 293T cell lines transfected with control siRNA or PLCE1+PLCB1 siRNA and cultured for one day. PLCE1 expression is not calculated for human zygotes due to the low expression level.

**Extended Data Figure 4:**
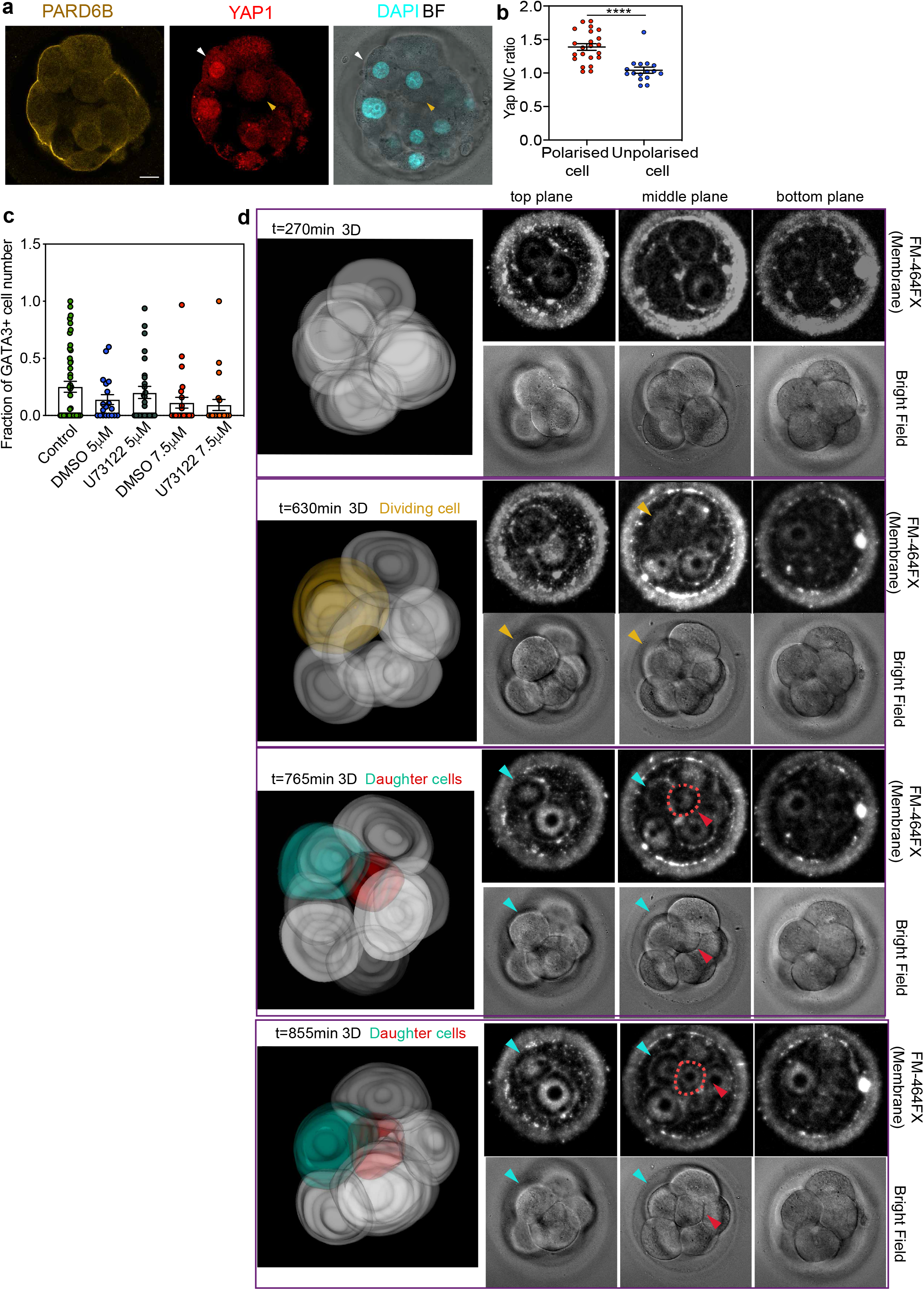
YAP1 display a higher Nucleus-to-cytoplasmic ratio in polarised cells at the morula stage. **a,** Human embryos at embryonic day 4 were immunostained for YAP1, PARD6 and DAPI. White arrows indicate outer cells, yellow arrows indicate inner cells. **b,** Quantification of Nucleus-to-cytoplasmic ratio of YAP1 in embryos from panel a. Each dot represents an analysed cell. N=22 polarised cells and N=18 unpolarised cells from 4 embryos analysed. ****p<0.0001, Mann-Whitney test. **c,** Quantification of fraction of GATA3+ positive cells for each embryo in different conditions shown in Fig. 3a. **d,** time-lapse imaging of cell position in compaction process during human embryo development. After a cell division, one of the two daughter cells was positioned to the inside of the embryo, and due to the compaction process this daughter cell was ultimately located to the inner position of the embryo. The dividing cell was labelled in yellow, and the two daughter cells were coloured with green and red. Red dotted lines indicate the red cell shown in 3D reconstruction. The red cell become localised to the inside. Scale bars, 15 μm.

## Material and Methods

### Ethical approval

This study was performed in four avenues: Clinical Embryology Laboratory at IVIRMA Valencia (Spain), University of Cambridge (United Kingdom), National Research Center for Assisted Reproductive Technology and Reproductive Genetics (China), and California Institute of Technology (United States).

The work in Clinical Embryology Laboratory at IVIRMA Valencia was approved by the National Commission of Human Reproduction (CNRHA), the General direction of research, innovation, technology and quality and by our institutional review board, the ethics committee of Clinical Research IVI Valencia, which complies with Spanish law on assisted reproductive technologies (14/2006). A total of 260 donated day 3 human embryos from 95 IVF patients were used. The average age of women was 28.12 ± 4. The fixed embryos were analysed at the University of Cambridge. This work was evaluated by the Human Biology Research Ethics Committee of the University of Cambridge (reference HBREC.2017.27).

The work performed at the University of Cambridge was in accordance with the Human Fertility and Embryology Authority (HFEA) regulations (license reference R0193). Ethical approval was obtained from the “Human Biology Research Ethics Committee” of the University of Cambridge (reference HBREC.2017.21). Informed consent was obtained from all patients from CARE Fertility Group and Herts & Essex fertility clinics, who donated surplus embryos used in this study after completing their IVF treatment. Prior to giving consent, patients were informed about the specific objectives of the project, and the conditions that apply within the license, offered counselling and did not receive any financial inducements. In this study, we used 15 donated embryos at the 2 pronuclei stage (day 1 d.p.f.), which were warmed and cultured in the Cambridge Laboratory according to the above regulations.

The work performed in the National Research Center for Assisted Reproductive Technology and Reproductive Genetics was conducted under the regulations of the Human Biomedical Research Ethics Guidelines (regulated by National Health Commission of the People’s Republic of China on 1 December 2016), the 2016 Guidelines for Stem Cell Research and Clinical Translation (issued by the International Society for Stem Cell Research, ISSCR) and the Human Embryonic Stem Cell Research Ethics Guidelines (regulated by China National Center for Biotechnology Development on 24 December 2003). These regulations and guidelines allow human gametes, and/or human embryos created or genetically manipulated *in vitro* and those cultured for no more than 14 days, to be used for scientific researches. The aim and protocols involved in this study were reviewed and approved by the Institutional Review Board of Reproductive Medicine, Shandong University. The protocols include the siRNA injection of human embryos.

The work at California Institute of Technology was approved by the California Institute of Technology Committee for the Protection of Human Subjects (Institutional Review Board number 19-0948). Funding was obtained through Open Philanthropy Project fund at the Silicon Valley Community Foundation. Human embryos at the zygote stage were obtained from the University of Southern California (USC) through the preexisting USC Institutional Review Board-approved Biospecimen Repository for Reproductive Research (HS-15-00859) after appropriate approval was obtained unanimously from the Biorepository Ethics Committee.

### Patient and embryo selection

In IVIRMA Valencia the majority of the day-3 embryos used in this study were generated by intracytoplasmic sperm injection (ICSI) (76.2%=198/260) using donor eggs (72.7%=189/260). The mean age of female patients providing oocytes was 28.12 ± 4 years (N=260), which is one of the strongest predictors of embryonic competence and oocyte quality^26^. The mean age of male patients providing sperm was 40.8 ± 6.98 years (N=260). The mean cell number at the time of embryo warming was 7.98 ± 1.26 (N=260), which is consistent with the 8 cells expected on day-3 stage of development of a good quality human embryo^27^. Most common female indications for Assisted Reproductive Technology included age (55%=143/260), poor ovarian response (8.1%=21/260) and tubal factor (7.7%=20/260).

All embryos used in National Research Center for Assisted Reproductive Technology and Reproductive Genetics were donated surplus 3PN zygotes generated from *in vitro* fertilisation, with the donor females age from 22 to 40 years old. The newly generated and donated 3PN embryos were used directly for microinjection experiments.

For embryos used in USC, all participants gave prior informed consent at USC Fertility to donate surplus cryopreserved embryos to research protocols after completion of childbearing or IVF treatment. Patients donated embryos of their own volition without coercion and did not receive any financial compensation for participation. A total of 41 donated human embryos at the zygote stage 2 pronuclei stage (day 1 d.p.f.) from 4 IVF patients were used. The average age of women was 44. The embryos were warmed at USC Fertility per usual IVF procedure and then transferred to the California Institute of Technology for the remainder of the protocol.

### Embryo warming and culture conditions

For experiments performed in Clinical Embryology Laboratory at IVIRMA Valencia: Vitrification and warming procedures were performed by means of the Kitazato method, as described elsewhere^28,29^. Briefly, embryos were equilibrated at room temperature in 7.5% (vol/vol) ethylene glycol + 7.5% dimethylsulfoxide (DMSO). After volume re-expansion, embryos were transferred to the vitrification solution consisting of 15% ethylene glycol + 15% DMSO + 0.5 M sucrose. After 1 min in this solution, embryos were placed on the Cryotop device using a minimum volume and were directly submerged in liquid nitrogen. For warming, the Cryotop was removed from liquid nitrogen and placed in 1.0 M sucrose in tissue culture media M 199 + 20% synthetic serum substitute (SSS) at 37°C. After 1 min, embryos were transferred to a solution containing 0.5 M sucrose at room temperature for 3 min. After two washes of 5 and 1 min, each in TCM199, embryos were cultured in a Geri Dish^®^ at 5.5 % CO2, 5% O2 and 37°C, humidified environment in a time lapse incubator Geri^®^ (Genea Biomedx, Australia). Embryos were cultured for 2, 9 or 24 h in pre-equilibrated Sydney IVF Blastocyst Medium (Cook, USA) or in the same medium supplemented with either U73122 - a PLC inhibitor (Caymanchem, USA, in 35 or 53mM DMSO) or DMSO (35 or 53mM, Sigma, USA) as vehicle control, without mineral oil. All culture media were pre-equilibrated to the incubator’s conditions overnight (5.5% CO2, 5% O2 and 37°C, humidified environment).

For the experiment performed at the University of Cambridge: Cryopreserved day 1 embryos were warmed using Origio thaw kit (REF10984010A)) following the manufacturer’s instructions. Briefly, the day before warming, Global Total human embryo culture medium (HGGT-030, LifeGlobal group) was incubated at 37°C + 5% CO_2_. Upon warming, the straw containing the embryo was defrosted at room temperature for 30 sec, and immersed in prewarmed (37°C) water for 1 min. The embryo was then transferred into vial 1 (5min), vial 2 (5 min), vial 3 (10 min) and finally in vial 4 for slow warming procedure. All these incubation steps were done using 12 well plates (ThermoFisher, 150628) and 1 ml per solution. Warmed embryos were finally incubated in drops of pre-equilibrated Global Total human embryo culture medium under mineral oil (9305, Irvine Scientific). Culture conditions are the following: 37°C 21% O_2_ and 5% CO_2_. Embryos were incubated for a total of 48 h in Global reaching 6-8 cell stage (day 3). At day 3 the embryos were either transferred into 96-well plate with each well containing 200 μl solution: control group (Global medium only), treated group (Global medium + PLC inhibitor at the concentration of 7.5μM, or to transfer to medium containing 35ng/ml FM 4-64FX for live-imaging. Embryos were stopped after 24 h culture via 4% PFA fixation.

For the experiment performed at California Institute of Technology, the embryos were warmed using Embryo Thaw Media Kit following the manufacturer’s instructions (Fujifilm Irvine Scientific, Cat. No. 90124). The day before warming, Continuous Single Culture-NX Complete medium (Fujifilm Irvine Scientific, Catalog No: 90168) was equilibrated overnight at 37°C + 5% CO_2_. On the day of warming (day 1), the straw that contains the embryo was defrosted at room temperature for 30 sec and immersed in prewarmed (37°C) water for 1 min until ice melted. The embryo was then transferred into T-1 (5 min), T-2 (5 min), T-3 (10 min) solutions for slow warming and finally into Multipurpose Handling Medium (MHM, Fujifilm Irvine Scientific, Cat. No. 90163) for recovery. All these incubation steps were done using 4 well plates (Nunc) and 1 ml per solution. Warmed embryos were finally incubated in drops of preequilibrated Continuous Single Culture-NX Complete medium under mineral oil (9305, Irvine Scientific). Culture conditions are the following: 37°C 21% O_2_ and 5% CO_2_. Embryos were incubated for a total of 48 h until reaching the morula stage (day 4). Embryos were fixed with 4% PFA at day 4 for immunofluorescence analysis.

### Experimental setting

A total of 60 day 3 embryos were cultured for 2 (N=21 embryos), 9 (N=26 embryos) and 24 h (N=13 embryos) after warming. Following fixation in 4% PFA, the compaction and polarisation timings of the human embryos were assessed by immunofluorescence analysis and time-lapse imaging evaluation.

To test the role of PLC, 8 experiments were performed with a total of 208 day 3 human embryos, which were warmed and randomly allocated into one of the following experimental groups: 5 μM U73122 (N=36), culture medium control (N=36) and vehicle control (N=23). Three additional series using increasing concentrations of U73122 were also performed: 7.5 μM U73122 (N=33), and its respective culture medium control (N=26) and vehicle control (N=25); and 10 μM U73122 (N=10), and its respective culture medium control (N=10) and vehicle control (N=9). All embryos were cultured for 24 h and fixed in 4% paraformaldehyde (PFA) to analyse cell polarisation and TE specification.

Embryo development was recorded in a Geri incubator (Genea Biomedx, Australia), which was programmed to acquire images of each embryo every 15 min through 11 different focal planes. The time-lapse videos of embryo development were analysed using the manual annotation software Geri Assess^®^ 1.0 (Genea Biomedx, Australia). The morphokinetic analysis was performed from the moment of embryo warming (Day 3) until embryo fixation at 2, 9 or 24 h after warming.

### Human embryo microinjection

For the siRNA microinjection experiment performed at the National Research Center for Assisted Reproductive Technology and Reproductive Genetics, in-vitro fertilised 3PN zygotes were injected and cultured in G1.5 medium (Vitrolife, cat. no.10128); for siRNA microinjection experiment performed at California Institute of Technology, the warmed embryos at 2PN stage were placed in MHM and microinjections were performed in MHM. Embryos were then cultured in drops of pre-equilibrated medium overlaid with mineral oil as described above.

The following siRNAs were used for microinjection: human PLCB1 siRNA #1 (CACACTACCAAGTATAATGAA) (Qiagen, Hs_PLCB1_4, SI00115521); PLCB1 siRNA #2 (CAGAGATGATCGGTCATATA) (Qiagen, Hs_PLCB1_6, SI02781184); PLCE1 siRNA#1(CAGGGTCTTGCCAGTCGACTA) (Qiagen, Hs_PLCE1_1, SI00115521); negative control siRNA (UUCUCCGAACGUGUCACGUdTdT) (Qiagen, 1022076). A 20μM concentration of siRNA solution was used for injection.

### Embryo fixation and Immunofluorescence

After culture, embryos were washed in PBS (Sigma, cat. no. D8537) and fixed in a freshly-prepared PBS solution containing 4% PFA (EMS, cat. no. 15710) for 20 min. For experiments done in IVIRMA Valencia, fixed embryos were washed twice in a PBS solution containing 0.1% Tween-20 (Sigma, cat. no. P9416) and immediately placed into a 0.5 ml PCR tube within an oil-PBS-oil interphase. Tubes were stored at 4°C were shipped to the University of Cambridge for immunofluorescence. For experiments done in other avenues, the embryos were directly processed for the subsequent steps.

A total of 203/266 embryos were subjected to immunofluorescence. Embryos were permeabilised in PBS containing 0.3% Triton X-100 and 0.1 M glycine for 20 min at room temperature. They were incubated in blocking buffer (PBS containing 10% BSA and 0.1% Tween) for 1 h. Primary antibodies were incubated overnight at 4°C in blocking buffer and secondary antibodies were incubated at room temperature for 2 h in blocking buffer. The following antibodies were used:

Primary antibodies:

Rabbit anti-PARD6(1:200)(Santa Cruz, sc-166405);
Goat anti-GATA3(1:200)(Thermo Fisher, MA1-028);
Mouse anti-aPKC(1:50)(Santa Cruz, sc-17781);
Mouse anti-YAP1(1:200) (Santa Cruz, sc-101199).

Secondary antibodies:

Alexa Fluor^®^ 568 Donkey anti-Goat;
Alexa Fluor^®^ 568 Donkey anti-Mouse;
Alexa Fluor^®^ 647 Donkey anti-Rabbit.
Alexa Fluor^®^ 647 Donkey anti-mouse.

In addition, embryos were stained with Alexa Fluor^®^ 488 Phalloidin (A12379, ThermoFisher Scientific) to reveal the F-actin cytoskeleton and with DAPI (D3571, ThermoFisher Scientific) to reveal nuclei.

Images were taken on an SP5 confocal microscope (Leica Microsystems) using a 40x oil objective N.A = 1.2. Laser power (less than 10%) and gain were kept consistent across different samples from the same experiment.

The images provided for siRNA injection experiments were taken on either an Andor Dragonfly spinning disc confocal microscope (63x oil objective N.A =1.4), or Sp8 confocal microscopy (Leica Microsystems) using a 40x oil objective N.A = 1.10.

### Cell culture, Transfection of small interfering (si) RNA with liposome

293 cells or HeLa cells were cultured in Dulbecco’s-modified Eagle’s medium (Invitrogen, cat. no. C11995500BT) added with 10% fetal bovine serum (FBS: Gibco cat. no. A3160801) and 1% penicillin-streptomycin solution (Gibco, cat. no.15140122) at 37¤C with 5% CO_2_. Transfections of siRNA were performed using Lipofectamine 3000 (Invitrogen, cat.no. L3000-015, Carlsbad, CA, USA) according to the manufacturer’s instructions. 293 cells or HeLa cells (1.6× 10^5) were plated onto 12-well plates and cultured until they were 60-70% confluent. After refreshing the medium, we mixed Lipofectamine 3000 and siRNA (Qiagen, Hs_PLCB1_4, SI00115521; Qiagen, Hs_PLCB1_6, SI02781184; Qiagen, Hs_PLCE1_1, SI00115521; Qiagen, negative control siRNA, 1022076) against target genes with Opti-MEM (Gibco) and the mixture of either control siRNA or PLCE1+PLCB1 siRNA were evenly added into each well. 293 cells or HeLa cells were further cultured for 24h to analyse the gene expression by qRT-PCR.

### RNA extraction and qRT-PCR validation

For 293 cells or Hela Cells, the RNeasy plus Micro Kit (Qiagen, cat. no.74034) was used for the extraction of total RNA following the manufacturer’s instructions. Genomic DNA was removed by digesting with RNase-free genomic DNA eraser buffer (Qiagen), and cDNA was obtained by reverse transcription of RNA using Evo M-MLV RT Kit for PCR Kit (Accurate Biotechnology(Human)Co.,LTD, cat. no. AG11603).

For human embryos, the zona pellucida was removed by mechanical dissection with a glass needle. Embryos were washed several times by gentle pipetting with a narrow-bore glass pipette to remove the attached cumulus or polar bodies. After final washing with 0.1% BSA/PBS three times, the embryos were collected and lysed in 2μl lysis buffer (0.2% Triton X-100 and 2 IU/μl RNase inhibitor) followed by reverse transcription with the SuperScript III reverse transcriptase. The PCR products were used for the templates of RT–PCR.

Power SYBR Green Master Mix (Takara) was used on a Roche 480 PCR system for qRT-PCR analysis. The mRNA level was calculated by normalising to the endogenous mRNA level of GAPDH (internal control) using Microsoft Excel. The qRT-PCR reactions were performed in triplicate for each experiment using gene-specific primers.The following primers have been used for qPCR: GAPDH: forward: GATCATCAGCAATGCCTCCT; reverse:TTCAGCTCAGGGATGACCTT; PLCB1: forward: GGAAGCGGCAAAAAGAAGCTC; reverse: CGTCGTCGTCACTTTCCGT; PLCE1: forward: TGCAGCCTCTCATCCAGTT; reverse: CCCTGCGGTAAATAGTCTGC.

### Data analysis

#### Time lapse imaging

Since the exact times of cryopreservation were not available in most cases, the following variables were assessed from the time of warming (t0) by means of time-lapse imaging: number of intact and degenerated cells, defined as the number of apparent viable and degenerated cells, respectively; number of total cells, defined as the sum of intact and degenerated cells; degree of fragmentation, defined as the estimated percentage of the embryonic volume occupied by cellular fragments; number of degenerating cells; defined as the number of cells degenerating during embryo culture; cell cycle length, defined as the time in hours between two consecutive cytokinesis in a given blastomere; time to the start of compaction, defined as the first time point after warming at which two adjacent blastomeres displayed an increased inter-blastomere angle; time to the end of compaction, defined as the first time point after warming at which all blastomeres displayed an increased interblastomere angle; duration of compaction, defined as the time interval between the start and end of compaction; number of cells at the start and end of compaction; and number of cells after embryo culture. Cell numbers shown in Extended Data Fig. 1 have been obtained using time-lapse images. To calculate cell numbers after compaction and after embryo culture, the number of cell divisions throughout the compaction stage and throughout the whole embryo culture were added to the number of cells before compaction and to the number of cells after warming, respectively. Hence, since some cell divisions may be untraceable when occurring inside the compacting embryonic mass, real cell numbers may be higher than reported.

All immunofluorescence images were analysed using Fiji^30^.

#### Apical enrichment analysis

F-actin and PARD6 polarisation were measured in a single focal plane, by taking the middle plane of the embryo. A freehand line of the width of 0.5μm was drawn along the cell-contact free surface (apical domain), or cell-contact (basal) area of the cell, signal intensity was obtained via the Region of Interest (ROI) function of Fiji. The apical/basal signal intensity ratio is calculated as: *I*(apical)/*I*(basal). A cell is defined as polarised when the ratio between the apical membrane and the cytoplasm signal intensity exceeds 1.5.

#### GATA3 expression analysis

the nucleus of each cells are masked using the Region of Interest (ROI) tool of Fiji. The average signal intensity of the ROI is calculated and a cell is defined as GATA3 positive when the nucleus to cytoplasm signal intensity exceeds 1.5.

#### Cell position analysis

embryos were imaged across their entire Z volume. By looking at F-actin localisation, the area of the cell of interest was examined on each plane. Cells that had an area exposed to the outside were defined as outer cells, whereas cells that did not have an area exposed to the outside were defined as inner cells.

#### Cell size measurements

we analysed the size of blastomeres in 8-cell stage human embryos (reference) and in human embryos harbouring inside cells. Cell size was measured in a representative single plane using the F-actin channel and manually drawing the contour of the cells.

#### Inter-blastomere angle

to measure the angle between two adjacent blastomeres we used the angle tool in Fiji and applied it to interfaces between blastomeres that were clearly visible in a given Z plane. 120° was chosen as a cut-off. The initiation of compaction was established as the time point at which one of the inter-blastomere angles in an embryo was above 120°. The end of completion of compaction was established as the time point at which all inter-blastomere angles in an embryo were above 120°.

#### 3D reconstruction and Segmentation

for 3D reconstruction images provided in Extended Fig. 4d, the membrane region of each cell at each plane was determined by FM-464FX signal, and the region of a cell across all planes were segmented manually using Fiji, the segmented masks were then transferred to 3D slicer software for the generation of 3D reconstructed images.

### Statistics and reproducibility

Statistical analyses were done using GraphPad Prism. Investigators were not blind to group allocation. Sample size was not pre-defined and embryos were randomly allocated into different experimental groups. Qualitative data was analysed using a Chi-square test. Quantitative data that presented a normal distribution was analysed using a two-tailed Student’s t test or an ANOVA test. Quantitative data that did not present a normal distribution was analysed using a Mann Whitney U test or a Kruskal Wallis test.

### Data availability

Data is available from the corresponding author upon request.

